# Decoding task representations that support generalization in hierarchical task

**DOI:** 10.1101/2024.12.02.626403

**Authors:** Woo-Tek Lee, Eliot Hazeltine, Jiefeng Jiang

## Abstract

Task knowledge can be encoded hierarchically such that complex tasks can be built by associating simpler tasks. This associative organization supports generalization to facilitate learning of related but novel complex tasks. To study how the brain implements generalization in hierarchical task learning, we trained human participants on two complex tasks that shared a simple task and tested them on novel complex tasks whose association could be inferred via the shared simple task. Behaviorally, we observed faster learning of the novel complex tasks than control tasks. Using electroencephalogram (EEG) data, we decoded constituent simple tasks when performing a complex task (i.e., EEG association effect). Crucially, the shared simple task, although not part of the novel complex task, could be reliably decoded from the novel complex task. This decoding strength was correlated with EEG association effect and behavioral generalization effect. The findings demonstrate how task learning can be accelerated by associative inference.

**Significance Statement:** Humans can generalize knowledge of existing tasks to accelerate the learning of new tasks. We hypothesize that this generalization is achieved by building complex tasks that associate simple (sub)tasks that can be reused. Using electroencephalogram (EEG) data, we showed that constituent simple tasks can be decoded from the EEG data of humans learning new complex tasks. Crucially, when participants represent complex tasks as associations between multiple simple tasks, the simple tasks can be decoded from the new complex task, even when they are not part of the new complex task. These findings demonstrate the importance of the reinstatement of simple tasks in task learning through generalization.

## Introduction

Humans have the remarkable ability to efficiently learn new complex and cognitively demanding tasks. To do this, we organize task representations hierarchically^1–4^, such that complex tasks can be represented by associating simpler ones. That is, new complex tasks can be learned efficiently by associating existing simple task representations and inferring their direct and indirect connections^5,6^. For example, a barista can learn how to make a ‘café latte’ by associating the existing skills of making espresso and steaming milk. In studies of human memory, researchers have found that associating multiple memory items enables us to adaptively adjust behavior by predicting future events^7^ or inferring novel information^8^. This associative feature of memory representations has been extended to incorporate the cognitive control parameters and task demands^5,6,9–13^ that support adaptive, goal-directed behavior. Although the lateral prefrontal cortex (lPFC) has been shown to be involved in representing task rules for goal-directed behavior^14^, little work has addressed how simple tasks can be associated and generalized for building of complex task representations.

This associative feature of task representations allows complex tasks to share components, which enables humans to generalize from previous experience and quickly adapt to new, related tasks^15^. For example, humans can extract structure from experiences and infer novel knowledge from the structure, a phenomenon sometimes also termed zero-shot learning^16–18^. Generalization is based on association, but it requires additional inferential processes as it allows for learning without direct association or experience. This generalization in task learning may be attributed to associative inference, which refers to the ability to infer novel, indirect associations from partially overlapping associations^9,11,19–26^. For example, the barista who is skilled at the ‘café latte’ task (espresso + milk) and the ‘café mocha’ task (espresso + chocolate) can infer how to make a hot chocolate via the ‘making espresso’ task shared by the ‘café latte’ and ‘café mocha’ tasks. In the laboratory, we reported that generalization via associative inference accelerates task learning and improves task performance^6^. Although generalization by associative inference plays an important role in task learning^3,6,9^, how it is implemented in the brain is understudied.

Two prominent theories of association and generalization in episodic memory offer insights into the processes supporting abstract task learning and generalization. Integrative encoding theory^21^ posits that repeated co-activation of items (e.g., A and C) will create an associated memory representation (e.g., AC). Moreover, integration can occur between associated item pairs that share an item (e.g., AB and BC), which leads to an integrated representation (e.g., ABC) that encompasses all three memory items^19,21,24^. The integrated representation will be retrieved if a subset of the items is present. For example, presentation of AC is expected to reinstate item B as well as items A and C. On the other hand, recurrent interaction theory^20^ proposes that an association between two memory items forms a conjunctive representation, and these conjunctive representations are connected to each of the constituent memory items. If a representation is activated (e.g., by sensory input), its connected representations will also be activated (though to a lesser extent). For example, if two conjunctions AB and BC share a memory item B, activating the conjunction AB will also activate memory items A and B. B will further activate conjunction BC, which in turn activates C. In the end, the coactivation of A and C will facilitate the formation of an indirect association of AC.

While these two major theories differ in terms of their proposed mechanisms of generalization, they share an important prediction about how the generalization occurs between associated memory representations via associative inference. When there are multiple associated item pairs that share a component (e.g., A-B and B-C association), activation of non-overlapping items (i.e. A and C) should reinstate the latent (i.e. not presented; B) item, because of the pattern completion process of integrated representation (integrative encoding), or feedback signal from conjunctive representations that are linked with activated items^20^.

Applying this idea to the context of task learning, we predict that the reinstatement of the latent task from the complex task is expected to be key for generalization, which relies on the associations between simple task representations. Moreover, both theories predict that stronger latent task reactivation will lead to more efficient learning, as it supports the inference of indirect associations between constituent task representations, via stronger activation of integrated representations^19,21,24^ or stronger reinstatement of constituent representations that are connected with latent representation^20^. However, while the reinstatement of memory *items* for generalization has been extensively studied, whether the same neural mechanism applies to *abstract task representations*, and if so, how the latent representations support efficient task learning, has not been directly tested. Here, we predict that the co-activation of simple task representations during complex task learning leads to association between simple tasks, and these associations can facilitate the new complex learning consisting of indirectly associated simple tasks. Based on this prediction, we hypothesize that the generalization will lead to reinstatement of latent task representation. Additionally, we hypothesize that stronger neural evidence of associative activation will predict stronger generalization.

To test these predictions, we leveraged a behavioral paradigm of generalization in task learning^6^. In this paradigm, subjects performed tasks that require to attend task-relevant feature(s) based on the cue(s). During performance, we obtained high temporal resolution and high-density electroencephalogram (EEG) data, to which we applied decoding and representational similarity analysis (RSA)^10,27^. To preview the results, the behavioral data revealed faster learning of complex tasks consisting of indirectly associated simple tasks compared to control tasks consisting of equally practiced simple tasks that were not indirectly associated. This result replicates the behavioral generalization effect reported in^6^ and suggests that participants learned how to attend task-relevant feature efficiently by optimizing cognitive control mechanism. We then observed robust decoding of constituent simple task representations from the complex task EEG data, which validates our analysis before investigating generalization effect in the EEG data. Critically, we observed in the complex task EEG data above-chance decoding of the latent task (e.g., B) that is reinstated by the indirect association (e.g., AC). We further found that the strength of the latent task representation in complex task EEG data is linked to both the strength of the constituent simple task representations in complex task EEG data and generalization performance. In sum, the results suggest that complex task learning results in a network representing associated information that enables zero-shot task learning, (e.g., generalization) via the reinstatement of the latent task. The strength of generalization is linked to the strength of the association.

## Results

### Experimental design

Participants (n = 40) first learned six simple tasks (coded as A-F), each of which was a feature-based attention task defined on one out of the six stimulus features (Fig. 1A). On each trial, participants judged whether a cued feature appeared in a forthcoming target stimulus (Fig. 1B). Next, during the training stage, participants learned four complex tasks, each consisting of two simple tasks. The complex tasks, denoted as the pair of constituent simple tasks (e.g., AB means a complex task consisting of simple tasks A and B), were presented on alternate blocks (odd blocks: AB, DE, even blocks: BC, EF; Fig. 1C). The complex tasks were divided into two sets, such that each set contained two complex tasks sharing one simple task (e.g., AB and BC). After the training stage, participants performed four new complex tasks (AC, DF, AD and CF) in the test stage (Fig. 1C). Two of the new complex tasks could be generalized via associative inference because both simple tasks were associated with the same simple task in practiced complex tasks (e.g., AC from AB and BC; Fig, 1D) and are hence termed generalizable tasks. In contrast, the other two new complex tasks (e.g., AD and CF) could not be generalized using associative inference, as the constituent simple tasks did not overlap during the training stage. These were termed control tasks. Within this behavioral framework, we defined task representation as cognitive processes involving task-relevant feature information and further posited that a major component of learning involves the improved application of cognitive control and associative mechanisms, leading to optimized feature-based selection and enhanced behavioral performance, based on previous research^6^. This design enabled us to investigate the behavioral generalization effect by comparing performance between generalizable and control tasks, while also examining the temporal dynamics of associative and generalization effects through EEG data.

**Figure 1.**
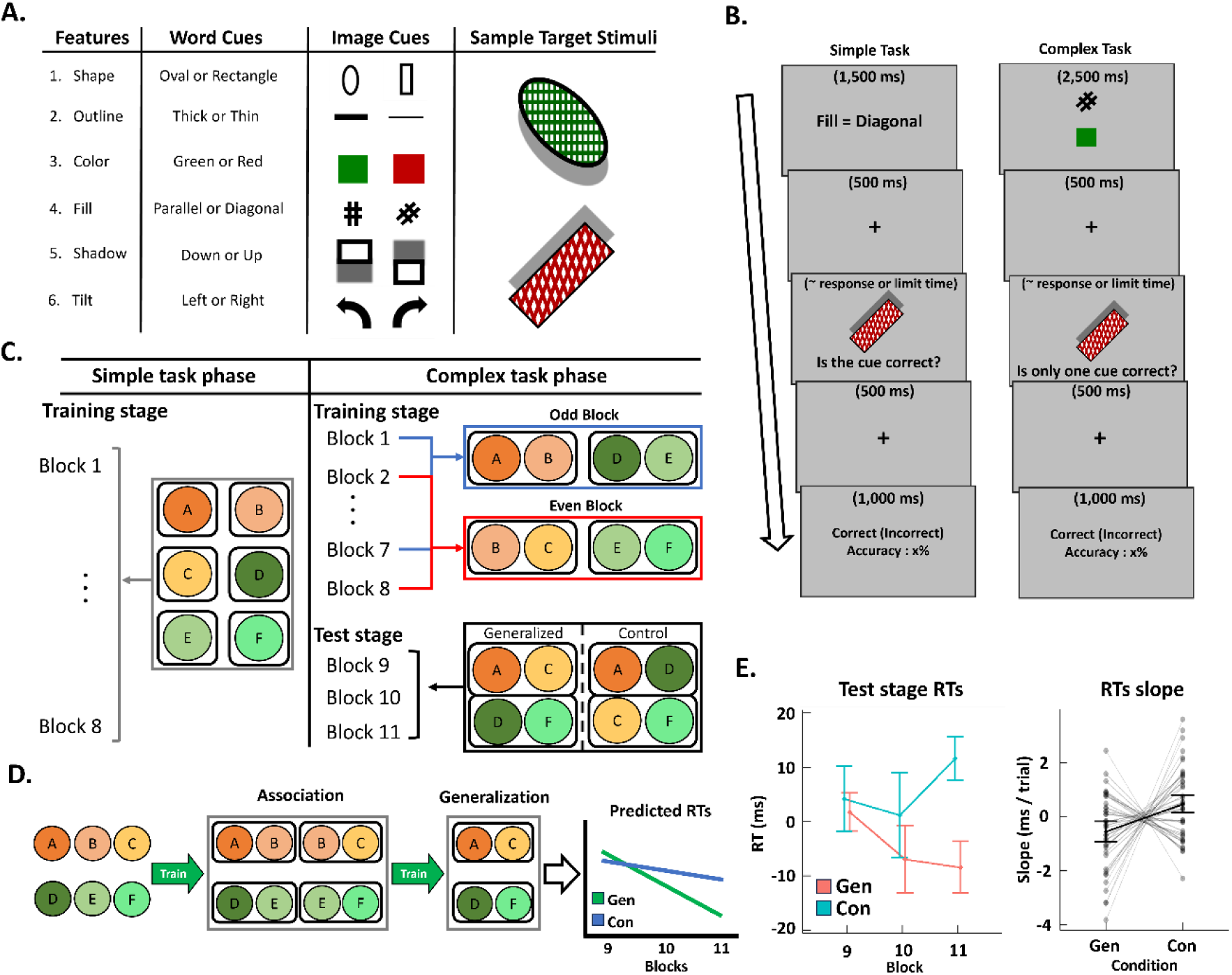
Experimental design and behavioral results. *(A)* Cue and target stimuli. Each cue indicated one stimulus feature and was presented as either text or image. Two example stimuli are illustrated in the rightmost column. *(B)* Trial events of the simple and complex tasks. In the simple task, participants first encountered a single cue that predicts a feature of an upcoming stimulus. After a delay of 500 ms, the target stimulus was presented until response or a deadline. Participants judged if the cue matched the target with button press. Visual feedback was provided after another 500 ms delay. The complex task is identical to the simple task except for two changes. First, participants encountered two cues instead of one. Second, participants decided whether only one cue matched the target (i.e., the ‘exclusive or’ rule). *(C)* Illustration of the experimental design. *(D)* Predicted dynamics of task representations in the complex task phase. Participants were expected to associate simple task representations to build a complex task representation and to further generalize two complex tasks via associative inference. Thus, we predicted that the generalizable complex tasks present faster decrements in RT than the control complex tasks. *(E)* Block-wise RTs (left, group mean and standard error of means, SEM) and slope of RTs (right, group mean and SEM, dots represent individual slopes) of complex tasks for each condition during the test stage.

### Behavioral data

Group-average accuracy for both simple and complex tasks stayed above 80% throughout the experiment (Fig. S1, S2, Note S1). Performance varied significantly across individual simple tasks, particularly for the ‘fill’ task (Table S1), so we conducted additional analyses to address potential confounding effects in the EEG data analysis driven by these behavioral differences, as detailed in the ‘Above-Chance Decoding in Simple Task EEG Data’ section.

To assess the generalization effect, we compared the performance of the generalizable and control conditions in the test stage (Table 1). Accuracy for the generalization and the control conditions did not differ significantly (t(39) = 1.56, p = .126, Cohen’s d = .25). For the RT analysis, we followed our previous procedure^6^ and regressed out confounding factors such as performance differences among simple tasks, response switch costs and so on (see Behavioral Analysis in Methods for details). The resulting residual RTs were used to test the hypothesis. Replicating our previous finding^6^, we found a significantly faster decrease in RT over time (t(39) = −2.35, p = .024, Cohen’s d = .37) in the generalizable complex tasks than in the control complex tasks (Fig. 1E), indicating faster learning for generalizable than control complex tasks.

**Table 1.** Behavioral results of the test stage in complex task phase. Group mean (SEM) of accuracy and residual RT in each condition (Gen = Generalizable, Con = Control) and block of the test stage.

| Condition | Block | Residual RTs (ms) | Accuracy (%) |
| --- | --- | --- | --- |
| Gen | 9 | 1.80 (3.51) | 78.54 (7.69) |
|  | 10 | -7.31 (6.17) | 79.17 (7.92) |
|  | 11 | -8.31 (4.75) | 81.77 (7.07) |
|  | Slope | -0.54 (0.23) per trial | 1.67 (1.11) per block |
| Con | 9 | 4.34 (6.03) | 74.69 (7.19) |
|  | 10 | 1.25 (7.77) | 79.48 (5.80) |
|  | 11 | 11.67 (4.00) | 79.17 (7.55) |
|  | Slope | 0.48 (0.22) per trial | 1.88 (1.11) per block |

While RTs for the control condition presented an unexpected incremental trend over blocks (Fig. 1E), this trend disappeared when we removed a regressor accounting for performance improvement over time in constituent simple tasks (Fig. S3). Without this regressor, we still found a marginally faster learning rate in generalizable condition than control condition (Fig. S3; t(39) = −1.99, p = .054). This result further supports our hypothesis by indicating that part of the learning effect in control condition could be attributed to the learning of constituent simple tasks, while generalizable condition showed significant benefit beyond the learning effect of constituent simple tasks. This benefit potentially involves the inference of indirect associations between simple tasks. This was tested using EEG decoding analysis and RSA presented below.

### Decoding simple task EEG data

For each participant and each time point (i.e., a 50 ms bin) of the simple task phase (Fig. 1C), we trained a multiclass decoder on the EEG data to classify the six simple tasks. The decoding analysis was performed after the onset of the target stimulus (Fig. 1B), so the decoder performance was not affected by perceptual processes related to the cue. Cross-validation decoding accuracy of simple tasks increased shortly after the target onset and was significantly above chance after 50 ms, peaking at around 300 ms (Fig. 2A, left panel). The longest epoch that showed significant decoding performance (indicated in red line in left panel of Fig. 2A) survived non-parametric permutation test for multiple-comparison correction (p_corrected_ < .001). The topographical map of the averaged pattern matrix^28^ shows strong absolute weights in occipital and posterior electrodes, with notable contributions from left frontal electrodes, suggesting both sensory and cognitive processes (Fig. 2A, right panel). This result suggests that the simple task representations can be robustly decoded in EEG data after target stimulus onset during their performance.

**Figure 2.**
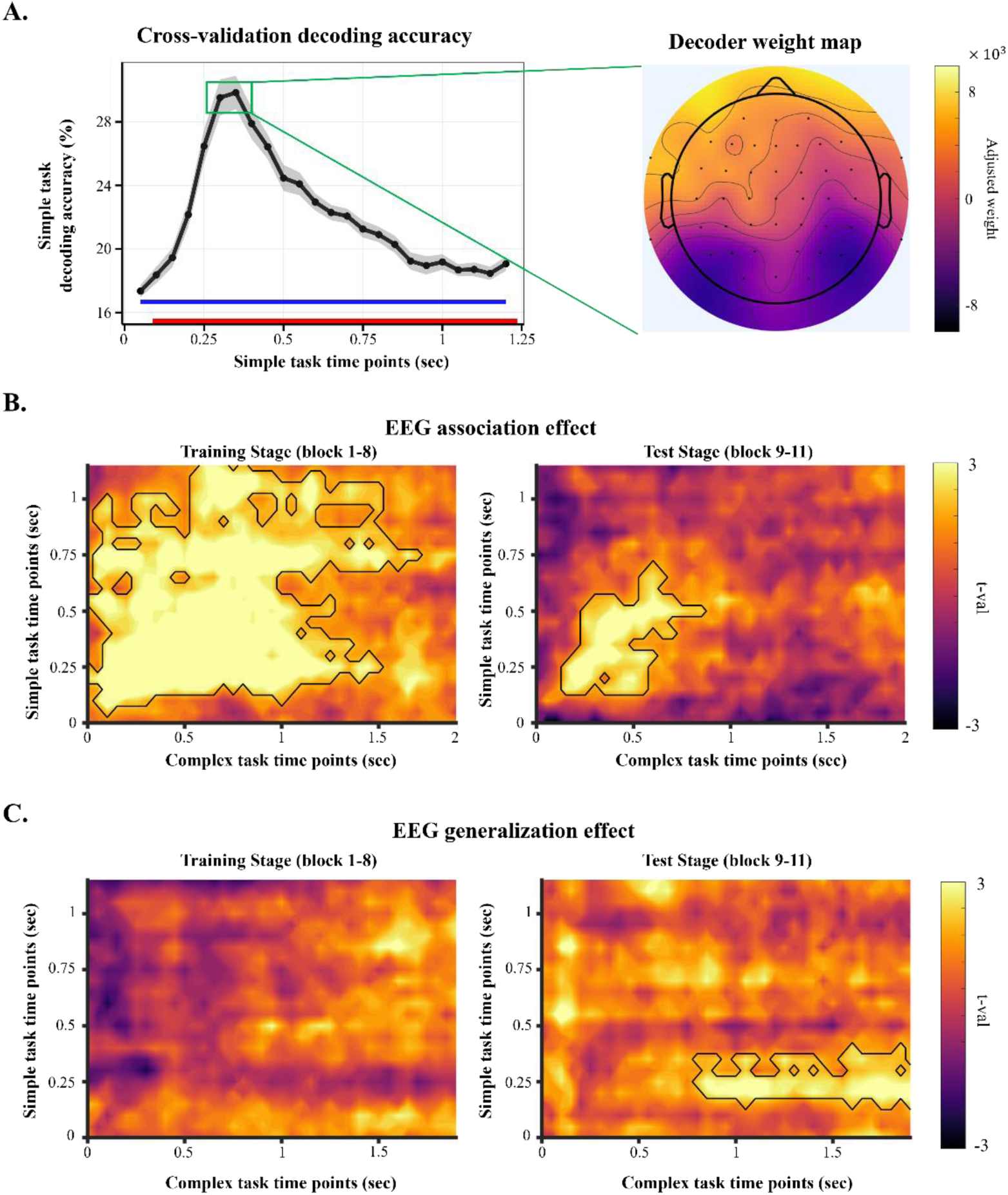
EEG decoding results. *(A)* Simple task EEG data decoding results. Left: Decoding accuracy over time within a trial. The grey shade indicates SEM of the decoding accuracy. The blue line indicates chance-level decoding accuracy. The red line indicates time points showing significantly above-chance decoding accuracy. Right: Visualization of mean electrode-wise contribution to the decoders based on pattern matrix^28^. *(B)-(C)* EEG association and generalization effects of complex task EEG data. In each panel, decoding performance (averaged t-values from trial-level decoding analysis, see Method) is plotted as a function of the time point of training (Y-axis) and the time point of test (X-axis). *(B)* Results of decoding constituent simple task representations from complex tasks in the training stage (left panel; e.g. A & B in AB) or in the test stage (right panel; e.g. A & C from AC). *(C)* Results of decoding the latent simple task in the training stage (left panel; e.g. C in AB), or in the test stage (right panel; e.g. B in AC). The time points are time-locked to the stimulus onset.

Due to the shared response deadline (see Methods) across simple tasks, participants showed significantly different behavioral accuracies between simple tasks (Table S1), especially low in the ‘fill’ task. To address the potential confounding effect in the decoder’s performance caused by the different number of trials across simple tasks, we calculated decoding accuracies separately for each simple task. Although the decoding accuracies differed among simple tasks, all tasks showed significantly higher decoding accuracies at most time points. More important, the decoding accuracies between simple tasks did not correlate with their behavioral accuracies, and ‘fill’ task showed similar, or higher, decoding accuracies compared to other simple tasks (fig. S4). This result indicates that the observed difference in behavioral performance between simple tasks did not systematically confound the decoders’ performance. But for further scrutiny, we included reaction times of each simple (or complex) tasks as a nuisance regressor in all decoding analyses and RSA.

### Overview of cross-phase EEG decoding analysis

To assess the reinstatement of simple tasks during the performance of complex tasks, we trained decoders on simple task phase EEG data, as described above, then tested on complex task phase data (see Method). To test the association effect (e.g., decoding simple tasks A and C from complex task AC), the decoders were tested on complex-task EEG data from the training stage (blocks 1-8). To test the generalization effect (e.g., decoding simple task B from complex task AC), the decoders were tested on the test stage of the complex task phase (blocks 9-11, Fig. 1C). For each test time point, the decoder produced a probabilistic prediction of each simple task being represented. The logit-transformed decoding probability was regressed against a linear model with two predictors representing composition and generalization effects separately (see Representational similarity analysis in Methods for more details). The trial-wise regression coefficient of each predictor was averaged for each participant and tested against 0 using a one-sample t-test. The presence of a composition/generalization effect predicts that beta values will be greater than 0. Thus, we used the t-values to quantify the strength of the composition and generalization effects. For the multiple comparison corrections, we applied a non-parametric permutation test^29^.

### EEG association effect

We first investigated the activation of constituent simple task representations to test the hypothesis that constituent simple task representations would be decodable during complex task trials. The purpose of this analysis was to assess the association between simple task representations for complex task learning from EEG data and to validate the simple task decoder’s performance when applied to the complex task EEG data. The decoding strength of constituent simple tasks can relate to the associations between task representations in two ways. One possibility is the subsequent memory effect^30–34^. In this study, subsequent memory effect posits that EEG decodability reflects neural representation strength of the constituent tasks, which in turn is linked to the encoding strength of the association between constituent tasks. Alternatively, associations between constituent tasks may drive stronger reinstatement in addition to the activation of the constituent tasks by the task demand of the complex task.

Figure 2B shows the time points that the constituent simple task representations can be decoded from the complex task EEG data (e.g., A & B from AB) in the training and test stages respectively. In the training stage, constituent simple task representations were decoded above chance level from complex task EEG data between 150 ms and around 1,200 ms post target onset when the decoders were trained between around 150 ms and 1,000 ms post target onset (p_corrected_ < .001; Fig. 2B, left panel). In the test stage, this EEG composition effect remained significant (p_corrected_ = .041; Fig. 2B, right panel) yet was reduced, significant between around 150 ms and 700 ms post target onset in training data and between 150 and 550 ms post target onset in test data. These results indicate that complex task execution involves constituent simple task activation, which is more extensive in the training than in the test stage.

### EEG generalization effect

While representing complex tasks as associations between simple task representations can be beneficial for their initial learning, generalization can support the learning of a novel complex task if its associations can be inferred from other learned complex tasks. Both the integrative encoding theory^21^ and the recurrent interaction theory^20^ predict the reinstatement of the latent task representation during test stage complex task trials (e.g., B from AC), despite the fact that the latent task is not present on the trial. To test this, we applied the decoding analysis and RSA to assess the reinstatement of the latent simple task representations in the complex task EEG data.

Consistent with the prediction, latent task representations were decoded above chance between 850ms to 2,000 ms post target onset in the test stage using decoders trained between 200 ms to 400 ms post target onset (Fig. 2C, right panel, p_corrected_ = .047). As the integrative encoding theory also predicts that the integrative memory can be formed after both partially overlapping associations are experienced^11,21^, generalization may occur in the training stage after both complex tasks from the same set were experienced (e.g., AB and AC). We also tested the decoding of the latent task in the training stage (e.g., C from AB). Note that the latent tasks in the training phase did not contribute to generalization via associative inference. In contrast to the test stage data, no time points reached statistical significance after the cluster-based non-parametric permutation test. These results suggest generalization between neural representations of complex tasks emerged mainly during the test stage, when complex task execution may benefit from generalization.

### Correlation between EEG association and generalization effects

Two major accounts of generalization in episodic memory^20,21^ hold that that the association effect (i.e., decodability of constituent simple tasks during complex task performance) lays the foundation of generalization effect (i.e. decodability of latent task from complex task in the test stage). Therefore, we predicted that the magnitude of the association effect should correlate with magnitude of the generalization effect. To test this correlation, we first calculated averaged EEG generalization effect for each participant within a time window defined in the following manner (Fig. 3A): Based on the EEG generalization effect in Fig. 2C, we trained decoders on simple task data from 200-300 ms post target onset. As the EEG generalization effect spans before and after responses, we divided the duration of the complex task showing the generalization effect (850- 2,000 ms post target onset, see Fig. 2C) into early and late parts using a threshold of the 75 percentiles of all complex task RTs across all participants (i.e., 1,100 ms). For consistency, the EEG association effect was trained on simple task EEG data from the same window (200-300 ms post target onset) and tested on time points showing statistically significant association effect, which starts at 100 ms (Fig. 2B, left). The test window was truncated at 850 ms to avoid temporal overlap between the periods used to measure the association and generalization effects.

**Figure 3.**
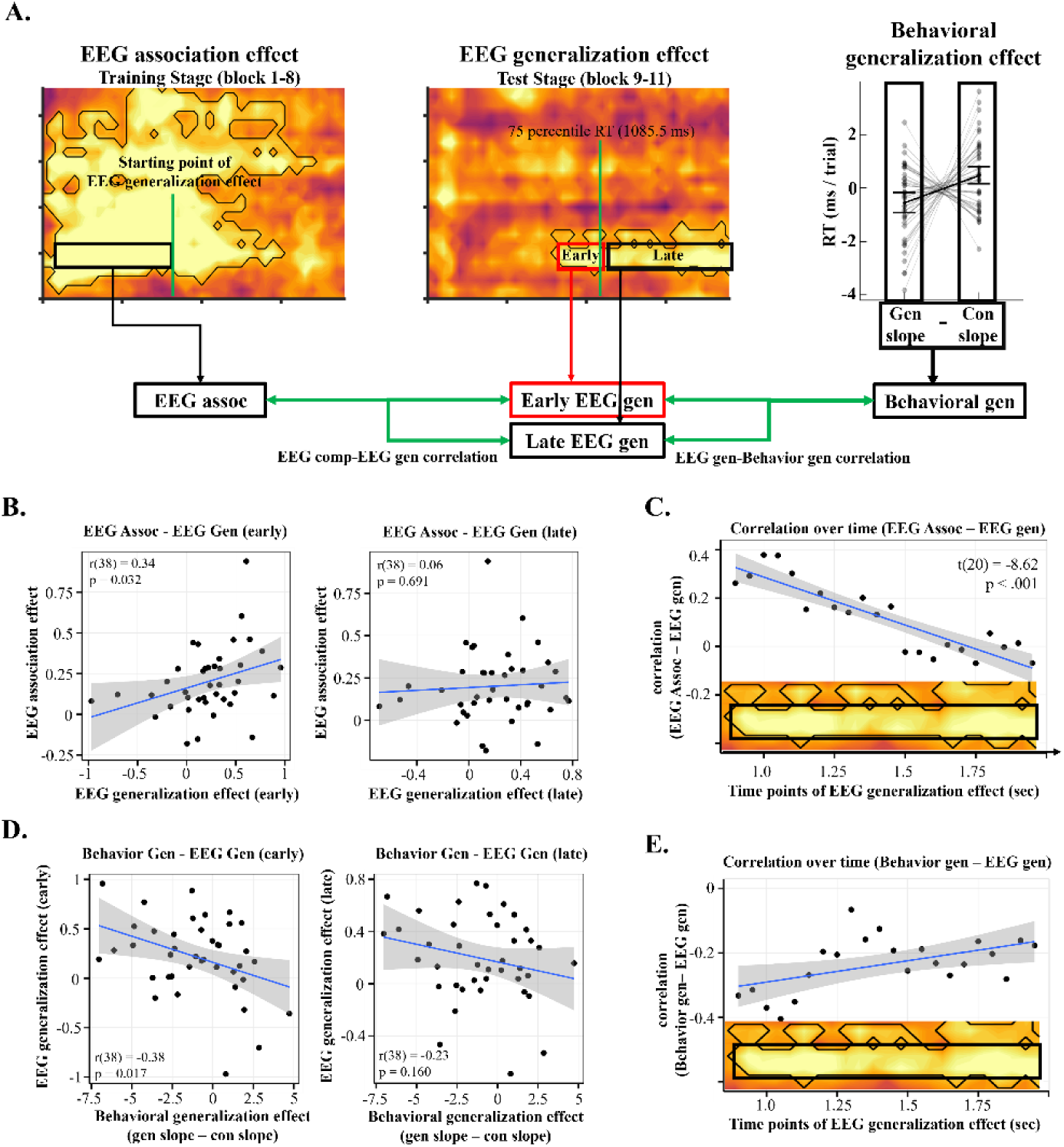
Correlation analysis results. *(A)* Data used for correlation analyses. For each individual, EEG association effect (left panel), EEG generalization effect (center panel), and behavioral generalization effect (right panel) were used. The EEG association effect was calculated by averaging single-trial t-values from the decoders trained on simple task time points between 200 ms to 300 ms and tested on training phase complex task time points between 100 ms and 850 ms. The EEG generalization effect was calculated by subject-mean single-trial t-values from the decoders to be trained on simple task time points between 200 ms to 300 ms and tested on complex task time points between 850 ms to 1,100 ms (early) or between 1,100 ms to 2,000 ms (late). The complex task time points were divided based on the 75% percentile RT (1085.5ms). The EEG association effect was calculated by subject-mean single-trial t-values from the decoders to be trained on simple task time points between 200 ms to 300 ms and tested on complex task time points between 100 ms to 850 ms. The behavioral generalization effect was calculated as the difference in RT slope (Fig. 1E) between generalizable and control conditions. *(B)* Scatter plots showing individual EEG association and early (left) or late (right) EEG generalization effects. *(C)* Correlation between overall EEG association effect and EEG generalization effect at each time point. *(D)* Scatter plots showing individual EEG generalization effect and early (left) or late (right) behavioral generalization effect. *(E)* correlation between behavioral and EEG generalization effects at each time point. The time points are time-locked to the onset of the stimulus.

Consistent with the prediction, EEG association effect in the training stage showed a significant positive correlation with the EEG generalization effect before the 75 percentile RT (r(38) = 0.34, p = .032) but not after (r(38) = 0.06, p = .691; Fig 3B). To rule out the possibility that the finding depended on the truncation of the time window, we repeated the analysis using the full time window (100-1,350ms). The untruncated EEG association effect exhibited a consistent pattern (before: r(38) = 0.39, p = .013; after: r(38) = 0.09, p = .589). To investigate the temporal dynamics of the correlation between association and generalization effects in the EEG data, we computed the average EEG generalization effect in the cluster (Fig. 3A, center panel) for each time point of the complex task and correlated it with the average EEG association effect. Note that this analysis did not rely on a threshold that segments trials into early vs. late conditions. We found a positive correlation between the EEG association effect and EEG generalization effect between 850 and 1,100 ms post stimulus onset. The correlation gradually attenuated and reached chance level at 1,500 ms. The correlation coefficient showed a significant decrease over time (Fig. 3C, slope: t(20) = −8.62, p < .001). Overall, these results indicate that the association of simple task representations during the training stage of the complex task phase was linked to the EEG generalization effect before and near the response in the test stage.

### Correlation between behavioral and neural effects of generalization

Lastly, we expected that the EEG generalization effect would benefit performance of generalizable complex tasks, whether it is via stronger activation of integrated memory representation with pattern completion process (integrative encoding), or via stronger activation of conjunctive representation that is connected with latent memory representation throughout training^20^. Therefore, we conducted exploratory analysis to investigate the behavioral relevance of the EEG generalization effect. We first assessed individual behavioral generalization effects using the RT slope difference between generalizable and control conditions (generalizable – control) and correlated this behavioral generalization effect with the EEG generalization effect from the previous analysis (Fig. 1E). The slope differences showed a significant negative correlation with the early part of the EEG generalization effect (r(38) = −0.38, p = .017), indicating that a stronger EEG generalization effect is linked to larger difference in RT decrease in generalizable tasks compared to control tasks. The correlation with the later part of the EEG generalization effect was not significant (r(38) = −0.23, p = .160).

To examine the temporal dynamics of the brain-behavioral correlation over time, we conducted the time-resolved correlation analysis above using EEG and behavioral generalization effect, similar to the previous section (fig. 3E). The change of correlation over time showed a significant positive linear trend (t(20) = 2.64, p = .016), indicating that in the test stage, participants who exhibited stronger EEG generalization effect before and near the response tended to show a stronger behavioral generalization effect.

## Discussion

We investigated the role of task representation reinstatement in supporting association and generalization in hierarchical task learning. To this end, we employed an established experimental paradigm that involves six simple tasks that can be combined to form complex tasks. Participants first trained with simple tasks, then learned complex tasks that consisted of two simple tasks. Specifically, participants learned complex tasks that shared a simple task (e.g., AB and BC) in the training stage. In the following test stage, participants performed generalizable complex tasks that consisted of indirectly associated simple tasks (e.g., AC) and control complex tasks that consisted of non-associated simple tasks (e.g., AD). Note that the generalizable complex tasks and control complex tasks are composed of constituent simple tasks that have received equivalent amounts of practice. Nonetheless, we hypothesized that participants would learn novel complex tasks efficiently by (1) associating learned simple task representations and (2) generalizing complex tasks via associative inference. Behaviorally, we successfully replicated the behavioral finding of greater slopes of decreases in RT for generalizable complex tasks than control complex tasks^6^, thus supporting generalization and association (as association is the foundation of generalization).

From the EEG data, simple task representations were reliably decoded from the complex task EEG data, supporting the association of simple tasks during the training stage, while it was weaker in the test stage. In our design, training directly co-activated constituent tasks, whereas in testing, reinstatement occurred indirectly via latent task representations. However, we cannot exclude the possibility that reduced reinstatement reflects weaker statistical power due to fewer trials in the test stage. These remain open questions for future research, as they highlight a limitation of our study in fully capturing the underlying mechanisms.

Critical for linking these data to generalization phenomena, we identified a significant latent task representation during the test stage but not during the training stage in complex task EEG data. Notably, this latent task was not required for new complex tasks, yet its reinstatement during the test stage supports the generalization of complex task representations. We also observed that a stronger EEG association effect during the training stage was associated with a stronger EEG generalization effect (measured prior to response) during the test stage. We further found that a stronger EEG generalization effect during the test stage at early time points of the trial was associated with a larger behavioral generalization effect. It is important to note that the task representation we decoded from the EEG data was not specific to the cue or stimulus, but rather cognitive processes involving task-relevant feature information, since the tasks required efficient guidance of feature-based attention. In addition, the decoder was trained with simple task EEG data to exclude potential confounding effects caused by the response paradigm (‘exclusive or’ design) from the complex tasks, which suggests that the decoded information is abstract task information.

EEG generalization effect suggests that the two major theories of associative memory regarding memory items can be further extended to apply to abstract task representations. Specifically, the latent task activation can be interpreted as a pattern completion process evoked by integrated representation that includes associated simple tasks^19,21,24^, or as an activation caused by link between conjunctive complex task representation and connected individual simple task representation^20^ (e.g., activation of A leads to an activation of AB, which further activates B). Both accounts are further supported by exploratory additional analysis using control complex task (Fig. S5), showing weaker indirectly linked task representations in similar time points. Overall, these results extend two major theories of associative memory, which focused on specific memory items, into task learning involving abstract representations.

The EEG association and generalization effects showed contrasting temporal dynamics in complex task trials. Constituent simple task representations appeared shortly after target onset (association effect, Fig. 2B), whereas latent task representations emerged around 900 ms later (generalization effect, Fig. 2C). Both associative episodic theories hypothesize that generalization relies on the reinstatement of the latent task, so the decoding of the latent task provides strong support for these theories. Integrative encoding theory^19,21,24^ suggests that the constituent simple tasks trigger pattern completion of the conjunction of all three tasks (e.g., ABC and DEF), leading to delayed activation of latent task^36^. On the other hand, recurrent interaction theory^20^ posits activation of a conjunction (e.g., complex tasks) will first activate the constituent simple tasks, which, after some delay, activate the latent task representation via recurrent connections. The observed temporal dynamics, where the EEG association effect precedes the generalization effect, align with the latter account. Nevertheless, future research is needed to adjudicate between the two theories in hierarchical task learning.

Our findings provide evidence for a neural mechanism underlying the flexible learning of complex tasks: compositionality. Compositionality has been hypothesized as a core process that enables efficient learning by leveraging existing representations to build higher-level ones^2,3,37,38^. In our experiment, the decoding analysis and RSA revealed reactivation of simple task representations during complex task learning, suggesting participants relied on them to build complex representations efficiently. This implies that compositional processing occurs during learning. However, while our results provide indirect evidence by demonstrating linear separability of task representations in higher-dimensional neural activity^37^, they do not directly confirm the compositional geometry of task representations. Future research should investigate this further by using recurrent neural network models, including state-space models^39^, which have demonstrated geometric trajectories of neural representations in multidimensional subspaces.

Consistent with generalization, we observed a significant correlation between the EEG association and generalization effect, suggesting that association may support generalization. Specifically, the early EEG generalization effect in the test stage correlated with the EEG association effect in the training stage, such that a stronger association of simple tasks during the training stage lead to stronger activation of the latent task representation during the test stage. Additionally, the relation between EEG association and generalization effects decreased over the time-course of the trial (Fig. 3B, C). We speculate that this pattern may be attributable to the different durations of latent task reinstatement across subjects. In other words, some participants may attenuate the reinstatement of the latent task sooner than others, which would compromise the correlation at later time points. Underlying the connection between the association and generalization effects are the associations linking the constituent simple tasks. In particular, when a simple task is shared by multiple complex tasks, it bridges multiple associations, which then form a network^20^. A remaining question is how to efficiently organize this network. One possible approach is to embed the network into a cognitive map, implying that spatial and non-spatial information can be organized into a map like space for goal directed behavior^40–44^. This would allow for efficient inference^41,45^. Another non-exclusive approach is representation compression^46^ that strengthens task associations to facilitate association and generalization.

In contrast to the zero-shot learning findings in previous studies^16–18^, our analyses showed a gradual improvement of performance in generalizable complex tasks. Although this may seem surprising, it can still be supported by the major theories of associative memories. Both posit that generalization via associative inference requires pre-existing association between memory items, as stronger association enhances pattern completion process^19,21,24^, or recurrent activation, leading to reinstatement of indirectly associated items. The late-stage reinstatement of latent task representations may provide a learning signal strengthen the associations between simple tasks, which are insufficiently formed during early training and become evident only later. Additionally, as the generalizable complex tasks were tested multiple times, this allowed for the effect of the learning signal to accumulate overtime. As a result, the generalization effect can be more manifested at the later stage of the test phase. Last but not least, while the previous research that reported zero-shot learning using generalization focused on the specific memory items relating strictly to declarative memory, our experiment required generalization of abstract task representations, which involves procedural memory that requires repeated practice for learning and is difficult to verbalize. These points may result in slow, gradual generalization processes, contrasting to the fast generalization effects that are reported previously^16–18^.

In conclusion, we provide electrophysiological evidence that the reinstatement of simple tasks supports complex task learning in hierarchical task organization. Specifically, we report the decoding of simple task representations from the complex task EEG data, which suggests that complex task representations include constituent simple task representations. Furthermore, we found significantly above-chance decoding of latent task representation from the EEG data of complex tasks that consisted of indirectly associated simple tasks, which suggests that participants generalized complex task associations via associative inference and the reinstatement of the latent simple task. Further supporting this conclusion, this latent task representation in the complex task EEG data was significantly related to the EEG measure of association effect and behavioral generalization effect. Together, these findings extend our understanding of hierarchical task organization that supports efficient task learning.

## Materials and Methods

### Participants

Forty-five healthy young adults participated in the experiment and received compensation at a rate of $25 per hour. Four participants were excluded from further analysis due to excessive non-stereotypical artifacts in the EEG data. One participant was excluded from the further analysis due to lower than 70% accuracy in a specific feature association during every block in training stages. The final sample consisted of forty participants (23 females and 17 males; age: M = 22.69, SD = 4.35). The experiment was approved by the University of Iowa Institutional Review Board.

### Stimuli

We used compound stimuli composed of six features. Each feature had two possible values (Fig. 1A). The features and their values are as follows: shape (oval or rectangle), color (green or red), outline (thick or thin), shadow (cast upward or downward), tilt (clockwise or counterclockwise), and pattern (parallel or diagonal). This design leads to 64 unique stimuli. Other stimuli included word and image cues indicating each of the feature values (Fig. 1A).

### Procedure

The experimental procedure was similar to Lee, Hazeltine and Jiang (6). The experiment consisted of two phases: a simple task phase (8 blocks of 48 trials each) followed by a complex task phase (11 blocks of 48 trials each). Participants had self-paced resting periods between blocks and had at least 5 minutes of rest between phases.

Each simple task trial started with the presentation of a cue of a stimulus feature value (e.g., filling pattern is diagonal) for 1,500 ms (Fig. 1B). The cue alternated between image and word format (Fig. 1A) after each trial so there were no cue repetitions. The cue was followed by the presentation of a fixation cross for 500 ms, after which the target stimulus was presented until response or a response deadline. The initial response deadline was set to 1,500 ms and was adjusted based on a staircase procedure. Specifically, the response deadline decreased by 50 ms when the participants made one or no errors and increased by 50 ms when participants made three or more errors in the last five trials. Otherwise, the response deadline remained unchanged. The participants were instructed to judge whether the cue matched the feature of the target stimulus by pressing the ‘D’ or ‘K’ button for yes or no, respectively. After each response, participants received feedback (correct or incorrect, presented in the center of the screen) for 500 ms. From the second block on, the feedback also included overall task accuracy. As a cue indicates which feature is task-relevant, the cues defined six different simple tasks (one for each cue). The simple tasks were coded using letters A-F (Fig. 1C), with the simple task-letter mapping randomized for each participant.

The complex task was similar to the simple task except for three modifications (Fig. 1B). First, the first screen showed two cues for 2,500 ms. Second, the rule of the complex task followed the ‘exclusive or’ rule. Specifically, participants were required to press the ‘D’ button if only one cue matched the target stimulus and the ‘K’ button otherwise. This rule was employed to ensure that the correct response could only be made after evaluating both cues. Third, the initial response time deadline was set to 2,500 ms for the staircase procedure. A complex task can be viewed as a combination of two simple tasks and was used to investigate association and generalization in hierarchical task representation. Thus, we coded a complex task using constituent simple tasks (e.g., complex task AB is composed of simple tasks A and B).

The complex task phase consisted of two different stages (Fig. 1C): a training stage (block 1–8) and a test stage (block 9-11). To study generalization at the complex task level, the simple tasks were divided into two groups (A-C and D-F). In the training stage, participants performed complex tasks AB and DE in odd runs and complex tasks BC and EF in even runs. In the test stage, participants performed generalizable complex tasks AC and DF, as well as non-generalizable control complex tasks AD and CF. Complex tasks AC and DF are generalizable because the constituent simple tasks were practiced with a common simple task (e.g., B and E) during the training stage. In contrast, control complex tasks AD and CF cannot be inferred by complex tasks learned in the training stage based on integrative encoding theory^19,21,24^ or the recurrent interaction of association theory^20^.

### Behavioral analysis

The behavioral analysis focused on replicating the generalization effect in Lee, Hazeltine and Jiang^6^. Specifically, we tested the generalization effect of faster learning of the generalizable than control complex tasks (Fig. 1D) using the analysis from Lee, Hazeltine and Jiang^6^. We first removed error trials and trials with RTs that were above 5 standard deviations from the subject mean. Next, we built a nuisance effect design matrix consisting of trial-wise status of response, response repetition, post-error, cue modality (text or image), task repetition, and cue modality repetition and regressed out the effects of these confounding factors from RTs in the test stage. Next, we fit a design matrix, which consisted of binary regressors marking generalizable and control complex tasks (to estimate RTs at the beginning of test stage) and their temporal order (to estimate RTs change over trials) against the residual RTs. These steps were performed separately for each participant. We compared the individual regression coefficients of estimated RT improvement over trials (slope) between regressors representing generalizable and control conditions of feature associations at the group level using paired t-tests.

### EEG recording & preprocessing

EEG data were recorded using a 64-channel active Ag/AgCl electrodes connected to an actiCHamp plus amplifier (BrainVision) at a rate of 500 Hz (10 s time-constant high-pass and 1,000 Hz low-pass hardware filters). The reference and ground were set to Pz and Fpz electrodes respectively. Raw EEG data was preprocessed using customized MATLAB scripts and the EEGLAB toolbox^50^. The EEG data were first band-pass filtered from 0.3 Hz to 50 Hz using Hamming windowed sinc FIR filter (pop_eegfiltnew.m). From the filtered continuous EEG data, we removed epochs with non-stereotypical artifacts for further analysis using outlier statistics and visual inspection. The remaining EEG data were re-referenced to common average and decomposed using the independent component analysis (ICA) to reject components reflecting eye movements and electrode specific artifacts. After excluding the artifact components, the EEG data was reassembled and epoched relative to the target stimulus onset (simple task: −1,000 to 1,250 ms, complex task: −1,000 to 2,000 ms) for further analysis (Fig. 4A). The choice of using target stimulus locked data was to control for the confounds of perceptual information when training and testing the decoders (see below). The preprocessed EEG data was smoothed and down-sampled to 20 Hz to reduce computation time for the decoding analysis and noise in the data.

**Figure 4.**
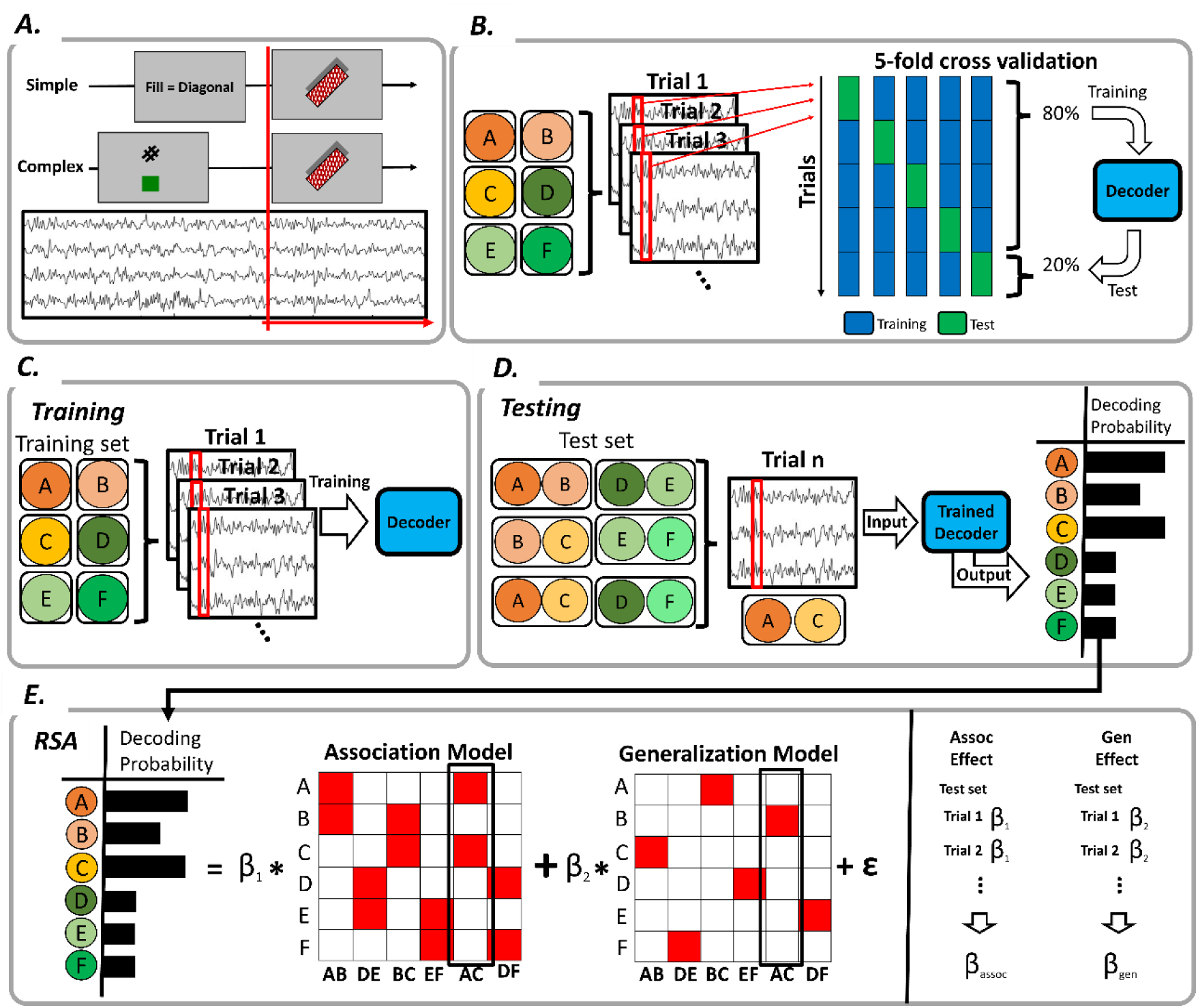
EEG decoding analysis procedure. (A) Illustration of the epochs for the EEG data analysis. The EEG epochs were time-locked to the onset of the target (red lines) to de-confound the effect of perceptual information on decoding performance. (B) Decoding analysis using simple task EEG data. At each time point, we trained and tested a decoder of six simple tasks using 5- fold cross-validation. (C) For cross-phase decoding analysis, the decoder was trained on simple task EEG data. (D) The trained decoder was tested using complex task EEG data and generated a decoding probability for each of the six simple tasks. (E) The RSA approach. The logit-transformed decoding probability was regressed against two model predictors representing association and generalization effects. This procedure was repeated on each complex task trial. We averaged the trial-wise association and generalization effects for each participant and then ran a one sample t-test against 0 for the statistical analysis.

### EEG decoding analysis

We trained decoders to discriminate among the six simple tasks at each time point using trial-level EEG data from all 64 electrodes (Fig. 4B). The decoders were L2 regularized multinomial linear regression with a tolerance (tol) of 1 × 10-4 and inverse of the regularization strength of 1^51^. Decoder training and test were implemented using the scikit-learn package^52^ and customized Python scripts. This approach was applied to 5-fold cross-validation using simple task data for each participant. The performance of the decoders was assessed by comparing the logit transformed probabilities to logit transformed baseline decoding performance (i.e., 1/6) using a one-sample t-test.

Following cross-validation, a decoder was trained on each time point using all simple task data (Fig. 4C) and tested on each time point and each trial of the complex task data ^53–56^ to assess the representation strength of simple tasks during the execution of complex tasks (Fig. 4D). As a result, for each test, the decoder generated a vector of six classification probabilities, each representing the decoder’s belief of a simple task being represented in the complex task at the time point of test. These probabilities were transformed with multinomial logit transformation for further analysis. The decoding results were stored in a four-dimensional tensor for each participant in the format of [trials in the complex task] × [time points of simple task] × [time points of complex task] × [6 simple tasks] for each trial of the complex task phase.

### Extracting pattern matrix from decoding analysis

Using the weight matrix (W) from the decoding analysis, we computed the pattern matrix (A) to quantify electrode involvement in decoding at each time point, ensuring better interpretability and biological relevance^28^. The calculation is given by:

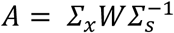

where W is the decoder’s raw weight matrix, *Σ*_*x*_ is the covariance of the EEG data across electrodes, and *Σ*_*s*_ is the covariance of the latent source estimates at each time point. This calculation generates a pattern matrix A that captures latent factors that are projected onto the electrodes. This method allows direct interpretation of the spatial contributions of each electrode to the decoded signal.

### Representational similarity analysis

For the RSA, we first tested association and generalization effects separately using the EEG data (Fig. 4E). The association effect is based on the hypothesis that neural representations of the complex task should include the encompassing simple tasks (e.g. AC = A + C). The generalization effect is built under the prediction that neural representations of the generalizable complex task should reinstate the latent task (e.g. simple task B is expected to be activated during complex task AC, if AB and BC are generalized). Each effect is represented as a predictor (i.e., a column in Fig. 4E). Next, the two predictors were included in a linear model, which was regressed against the logit-transformed classification probabilities from each trial of the complex task EEG data. From the output of linear regression, we acquired trial-wise t-value for each predictor, which represents the association/generalization effect on the current trial. For each regressor, the t-values were grouped as a vector with the length of trials in the complex task. In this vector, trials that showed extreme t-values (± 5 SD) were excluded following a previous protocol^10^. Each vector from the association and generalization effects was averaged to examine the overall fitness of the regressor across trials. The regression was conducted on decoding probabilities obtained when the decoder was trained on each time point of the simple task EEG data and tested on each time point of the complex task EEG data. In the end, these analyses generated a t-value matrix with the size of the number of time points of simple task × the number of time points of complex task for each participant. The statistical significance was calculated by conducting one sample t-test across participants against 0 at each of combinations of simple task and complex task time points.

### Cluster-based permutation test

To control for multiple comparisons in the above the final t-value matrix, we conducted a cluster-based permutation test with customized python script based on Maris and Oostenveld ^29^. Specifically, with simple task EEG data, we repeated a 5-fold cross-validation decoding analysis using shuffled labels of the simple tasks 10,000 times. We pooled the clusters with the largest sum of absolute t-values (t-sum) from each repetition to create a null distribution of t-sums, which takes into consideration of both the cluster size and statistical significance. From the actual result, the cluster with the largest t-sum is compared to the null distribution to obtain corrected p-value. For the RSA, we repeated the analysis using shuffled labels of the simple tasks before training the decoder 10,000 times while keeping the labels of the complex tasks the same between repetitions. The remaining procedure was identical to the non-parametric permutation test for simple task decoding analysis.

## Data availability

All data and analysis scripts related to this paper are available in Open Science Framework at (https://osf.io/mzf4a/).

## Acknowledgments

This project was supported by the National Institute of Mental Health (R01MH131559 to J.J.). We thank the members of the Cognitive Control Collaborative and the members of the Human Perception and Performance Group at the University of Iowa for helpful discussions.

## Author Contributions

**Woo-Tek Lee**: Conceptualization, Methodology, Investigation, Visualization, Writing—original draft, Writing—review & editing. **Eliot Hazeltine**: Conceptualization, Supervision, Writing—review & editing. **Jiefeng Jiang**: Conceptualization, Methodology, Investigation, Supervision, Writing—review & editing, Funding acquisition.

## Classification

Biological Sciences, Psychological and Cognitive Sciences.

