## Supplemental information for "Decoding task representations that support generalization in hierarchical task"

### Supporting Information for
Decoding task representations that support generalization in hierarchical task.

Woo-Tek Lee, Eliot Hazeltine, and Jiefeng Jiang

Corresponding author: Jiefeng Jiang


**This PDF file includes:**

Supporting text
Figures S1 to S5
Table S1

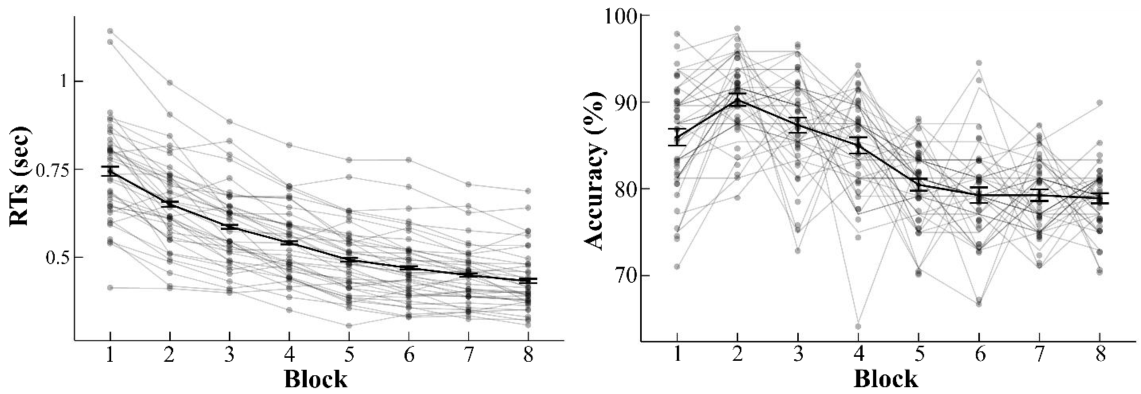


**Fig. S1.** Behavioral results from the simple task phase. Individual RT (left) and accuracy (right), imposed with the group mean and SEM, are plotted as a function of block. Participants showed faster RTs over blocks on correct trials, while the accuracy declined after block 2. The decline in accuracy may be attributed to the decrement of the response deadline due to the staircase procedure of adjusting response deadline.

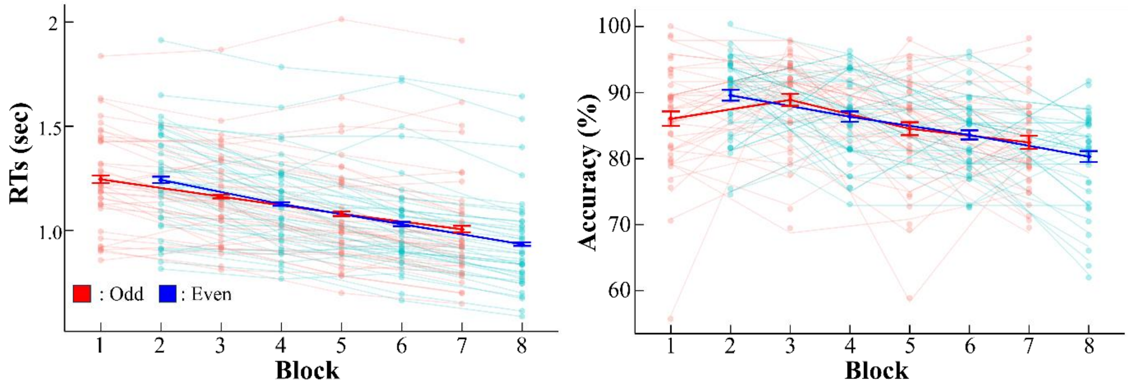


**Fig. S2.** Behavioral results from the training stage of complex task phase. Individual RT (left) and accuracy (right), imposed with the group mean and SEM, are plotted as a function of block. Participants showed faster RTs over blocks on correct trials, while the accuracy declined after block 3. The decline in accuracy may be attributed to the earlier response deadline due to the staircase procedure of adjusting response deadline.

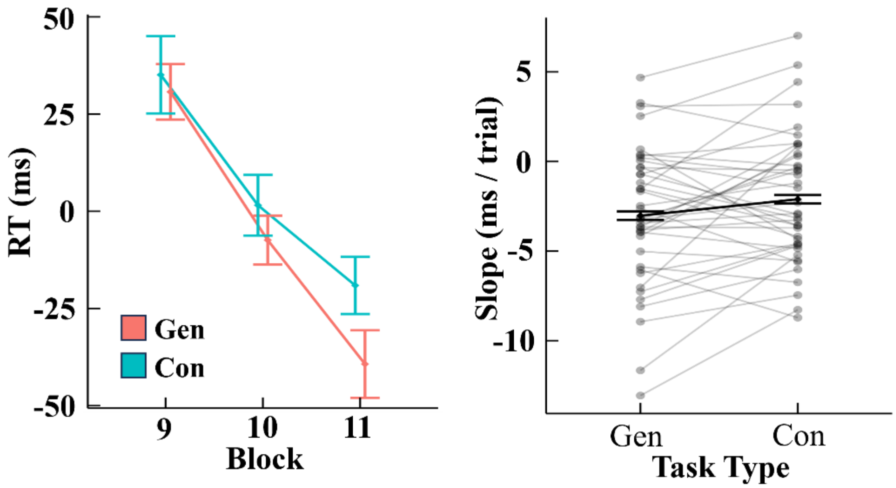


**Fig. S3.** Block-wise RTs (left, group mean and standard error of means, SEM) and slope of RTs (right, group mean and SEM, dots represent individual slopes) of complex tasks for each condition during the test stage, after removing a confounding regressor accounting learning effect of constituent simple tasks. RT slope showed marginally significant difference between generalizable and control complex tasks ($t(39) = -1.99$, $p = .054$).

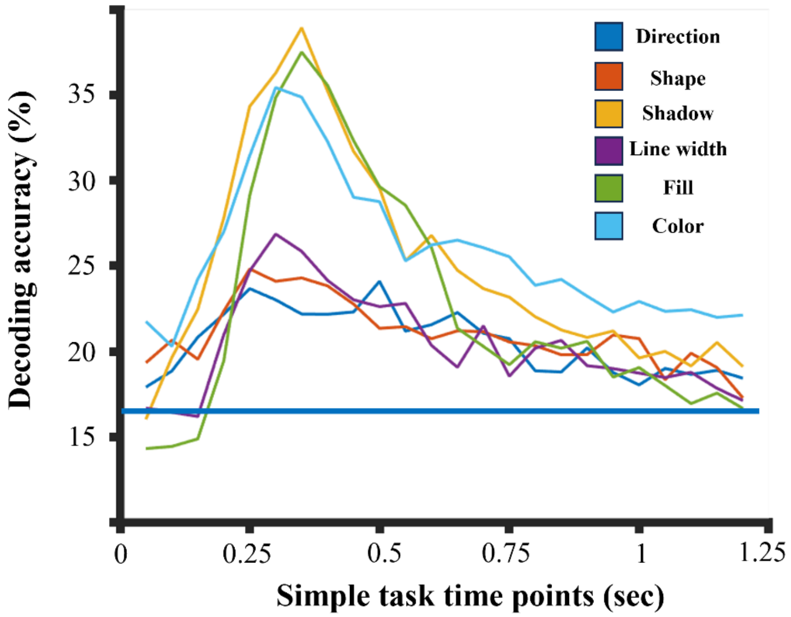


**Fig. S4** Simple task EEG data decoding results for each task. Blue flat line indicates chance-level decoding accuracy.

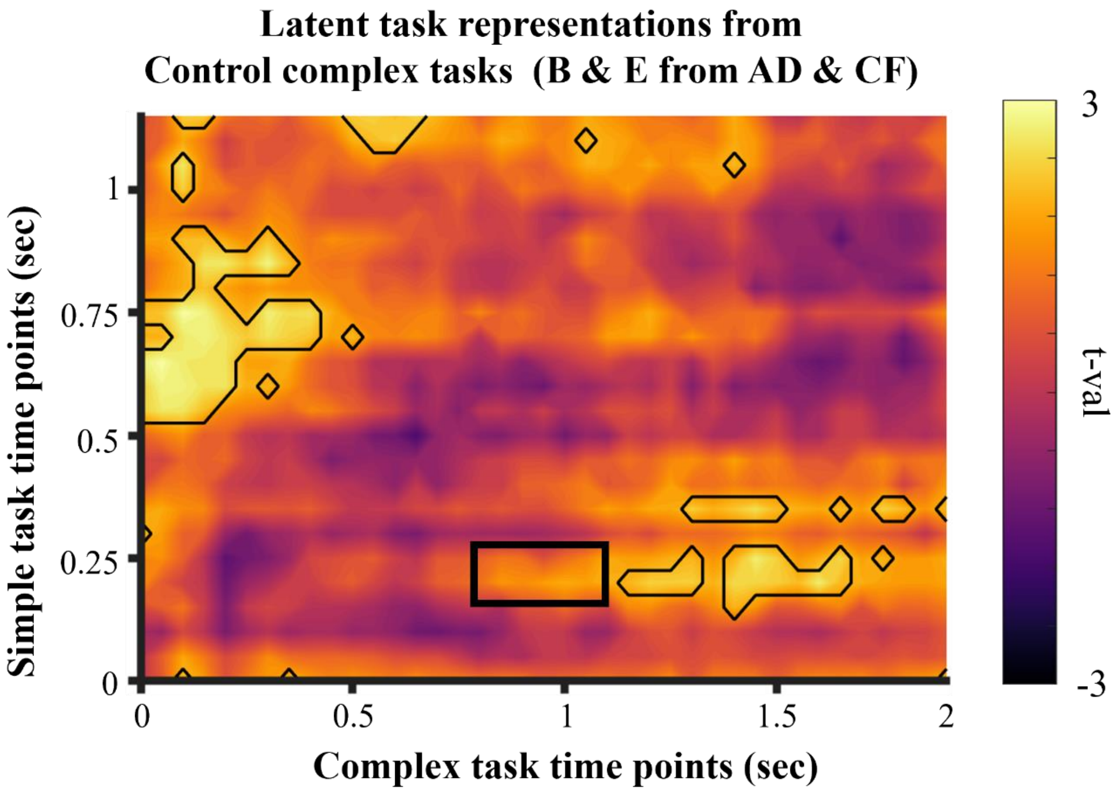


**Fig. S5** Result of decoding linked latent task representations (e.g., B & E from both AD and CF). These results are not applied cluster-based permutation tests. This result provides preliminary evidence that in control complex tasks the latent tasks are also reinstated, as predicted by the two major associative memory theories. The reinstatement was weaker than generalizable complex tasks, possibly because the latent tasks in control complex tasks were retrieved by one cue (e.g., B is only retrieved by A in complex task AD), whereas the latent task in generalizable complex tasks was retrieved by two cues (e.g., B is associated with both A and C in AC). An additional correlation analysis between the EEG generalization effect in control complex tasks, which were identical to the generalizable complex tasks (marked as a black box), and the behavioral generalization effect showed a marginal negative relationship ($r(38) = -0.29$, $p = .069$), suggesting that stronger reactivation of the latent task is associated with stronger behavioral generalization effect.

**Table S1**

Behavioral performance of each simple task in the simple task phase.

| | Tilt Direction | Shape | Shadow | Line Width | Fill | Color |
|---|---|---|---|---|---|---|
| Acc (%) | 86.3 (0.87) | 90.6 (0.67) | 84.2 (0.95) | 80.6 (1.15) | 64.1 (1.20) | 94.1 (0.56) |
| RTs (ms) | 535.5 (18.29) | 520.5 (15.95) | 577.1 (15.47) | 574.7 (16.77) | 652.9 (17.24) | 478.9 (13.70) |

*Note*. Values are reported as the ‘mean (standard error)’.

#### Supporting Information Text

**Linear mixed-effect model analysis.**

To test the learning effect in the training stage of complex task, we performed linear mixed-effect model analysis. The analysis was performed on RTs and accuracy separately using R software version 4.1.0 (1) with lme4 package (2). The formula for this analysis is as follows:

$$y_i = (\beta_0 + u_{i1}) + (\beta_1 + u_{i2})x + \varepsilon_i$$

Where $y_i$ refers to a value of behavior measures (either block-mean RT or accuracy) for a participant $i$. $\beta_0$ and $\beta_1$ each represent fixed effects for the intercept ($\beta_0$) and slope ($\beta_1$) that are consistent among participants (i.e., fixed effects). Additionally, $u_{i1}$ and $u_{i2}$ represent random effects for the intercept and slope that can vary for each participant. The term $x$ represents block, and $\varepsilon_i$ reflects errors. In this model, the effect of interest was the fixed-effect of slope, $\beta_1$, that indicates how the behavioral measure changes over blocks.

In the simple task, we found that RTs consistently decreased over blocks ($\beta_1$ = -0.04 (0.002), $t_{39}$ I= -20.45, p < .001, Cohen's d = -3.27). Accuracy exhibited a significant, yet small decline over time ($\beta_1$ = -1.58 (0.153), $t_{39}$ = -10.33, p < .001, Cohen's d = -1.65). The decline in accuracy likely reflects a speed-accuracy tradeoff resulting from the staircase procedure constantly reducing the response deadline (see Methods). Specifically, the staircase procedure will increase the difficulty by reducing the response deadline if the accuracy was less than 80% in the last five trials. Nevertheless, group average accuracy remained above 80% in all blocks. In the training stage of complex task, we found a pattern identical to the simple task for both RTs (Odd: $\beta_1$ = -0.04 (0.005), t39 = -7.53, p < .001, Cohen's d = -1.21; Even: $\beta_1$ = -0.05 (0.004), $t_{39}$ = -13.39, p < .001, Cohen's d = -2.14) and accuracy (Odd: $\beta_1$ = -0.01 (0.003), $t_{39}$ = -2.48, p = .017, Cohen's d = -0.40; Even: $\beta_1$ = -0.02 (0.002), $t_{39}$ = -7.61, p < .001, Cohen's d = -1.22) while maintaining a block-wise average accuracy of over 80%. Similar to the simple task phase, the decrease in both RT and accuracy is likely a result of speed-accuracy tradeoff as the response deadline shortened. Overall, the behavioral data indicate that the participants performed the simple and complex tasks as instructed.
